# Proteostasis in ice: The role of heat shock proteins and ubiquitin in the freeze tolerance of the intertidal mussel, *Mytilus trossulus*

**DOI:** 10.1101/2022.02.03.478032

**Authors:** Lauren T. Gill, Jessica R. Kennedy, Katie E. Marshall

## Abstract

The bay mussel, *Mytilus trossulus*, is one of the few animals that can survive internal ice formation. Freeze tolerant intertidal animals, like *M. trossulus*, may freeze and thaw many times during the winter, depending on air and ocean temperatures. Freezing can cause protein denaturation, leading to an induction of the heat shock response with expression of proteins like HSP70, and an increase in ubiquitin conjugated proteins. There has been little work on the mechanisms of freeze tolerance in intertidal species, limiting our understanding of this survival strategy. Additionally, this limited research has focused solely on the effects of single freezing events, but the act of repeatedly crossing the freezing threshold may present novel physiological or biochemical stressors that have yet to be discovered. We predicted that repeated freeze exposures would increase mortality, upregulate HSP70 expression, and increase ubiquitin conjugates in mussels, relative to single, prolonged freeze exposures. *Mytilus trossulus* from Vancouver, Canada were repeatedly frozen for a combination of 1 × 8 hours, 4 × 2 hours, or 2 × 4 hours. We then compared mortality, HSP70 expression, and ubiquitin quantity across experimental groups. We found a single 8-hour freeze caused significantly more mortality than repeated freeze-thaw cycles. We also found that HSP70 and ubiquitin expression was upregulated exclusively after freeze-thaw cycles, suggesting that freeze-thaw cycles offer a period of damage repair between freezes. This indicates that freeze-thaw cycles, which happen naturally in the intertidal, are crucial for *M. trossulus* survival in sub-zero temperatures.

## Introduction

Animals living in the intertidal zone – the area where the ocean meets the land between high and low tides – must contend with both marine and terrestrial conditions. Intertidal animals must survive intense UV radiation, hydrodynamic forces, desiccation stress, and extreme fluctuations in temperature, salinity, and pH (Carrington et al., 2009; Jensen & Denny, 2016; Pulgar et al., 2017). Thermal extremes are an especially important abiotic factor on rocky shores, determining both distribution and physiological performance of organisms (Helmuth & Hofmann, 2001). During the winter, sub-zero temperatures often induce ice formation in intertidal animals due to the constant wet conditions of the intertidal zone and the presence of food particles in their bodies which can act as ice nucleators (Storey & Storey, 1996). However, ice formation in intertidal ectotherms can inflict cellular and protein damage (Aarset, 1982). Despite this, intertidal invertebrates inhabit virtually all rocky coastlines in temperate and polar regions, and many are freeze tolerant, allowing them to survive the frigid conditions that occur during the winter months. Although there is a growing body of literature focusing on thermal tolerances of intertidal organisms, this literature focuses primarily on tolerance to heat stress, meaning that we are just beginning to understand the physiological mechanisms behind survival at low temperatures and how these mechanisms drive ecology.

Investigations into freeze tolerance of intertidal animals are further complicated by the fact that animals will often experience multiple freeze-thaw cycles throughout the winter. Repeated freezing in the intertidal zone is caused primarily by cyclic tidal regimes. When animals are immersed in sea water during winter, they are protected from freezing air temperatures and experience more moderate conditions (Kennedy et al., 2020). Therefore, sub-zero air temperatures can cause freeze tolerant organisms to freeze when the tide recedes, and then thaw upon immersion as the tide returns. This variation also means that organisms that are located higher in the intertidal zone spend longer amounts of time out of water and thus must tolerate more extreme abiotic pressures and more severe sub-zero body temperatures during daily tide cycles in the winter, as compared to animals located lower down on the shore (Davenport & Davenport, 2005; Finke et al., 2007; Williams & Somero, 1996). This time spent out of water means a cessation of feeding in suspension feeding animals, a decrease in metabolic rate, increased chance of desiccation, and other pressures (Storey & Storey, 1988). This suggests that since high intertidal animals will be exposed to freezing conditions for longer, they may be better acclimated to freeze exposures, as compared to their low intertidal counterparts.

In order to survive freezing, intertidal animals must be able to survive the formation of ice and then regain natural function once thawed (Storey & Storey, 1988). However, irrespective of ice formation, animals exposed to low temperatures can still experience considerable damage from the cold (Ramløv, 2000). Low temperatures can cause a decrease in cellular reaction rates, a suppression of protein synthesis, and promote destabilizing, hydrophobic interactions among protein residues rendering them non-functional (Al-Fageeh & Smales, 2006; Privalov, 1990; Ramløv, 2000). In addition, when exposed to sufficiently cold temperatures, internal ice formation can occur imposing mechanical and osmotic stress on the organism (Ramløv, 2000). As extracellular water is formed into ice, solutes are excluded, causing the surrounding fluids to become increasingly solute rich (Lee, 2010), which in turn, may lead to cellular protein denaturation (Baust, 1973). Finally, once the frozen animal begins to thaw and recover, damaging reactive oxygen species are generated, and potential further mechanical damage can occur due to ice recrystallization (Doelling et al., 2014; Mazur, 2010). Therefore, three distinct processes will impose challenges on intertidal organisms during freeze-thaw cycles: cooling, freezing, and thawing (Toxopeus & Sinclair, 2018).

To withstand damage due to freezing, animals can accumulate cryoprotectants, which function to protect against the effects of low temperature and ice formation (Storey & Storey, 2013). One such cryoprotectant, heat shock proteins (HSPs), are an important way that organisms can reduce the impacts of protein denaturation due to freezing. Heat shock protein 70 (HSP70) is a highly conserved, 70 kDa chaperone protein that acts as a chaperone to ensure cellular proteins are folded correctly in their native state (Boroda et al., 2020). In an environment with cold-induced protein denaturation, HSP70 also plays a role in re-folding misfolded proteins to regain their native states and can also target denatured proteins to remove them from the cell (Todgham et al., 2007). This removal can prevent the irreversible formation of cytotoxic protein aggregates (Venetianer et al., 1994).

Most previous work on HSP70’s role in an organism’s thermal tolerance has focused on upper thermal limits and heat shocks. Increased expression of HSP70 as a result of elevated temperature stress has been widely observed in intertidal organisms, such as *Mytilus galloprovincialis* mussels, *Lottia gigantea* limpets, and snail species of the genus *Tegula* (Miller et al., 2009; Tomanek & Sanford, 2003; Toyohara et al., 2005). However, much less is known about the extent of HSP70’s role in relation to cold stress. HSP70 is upregulated in *Epiblema scudderiana* moth larvae in response to winter declines in temperature, and plays a significant role in their winter survival (Zhang et al., 2018). HSP70 expression is also upregulated in fruit flies in response to acute cold shocks (Štětina et al. 2015). Similarly, larvae of the Antarctic midge*, Belgica antarctica,* expressed increased levels of HSP70 after freezing, indicating more protein damage (Teets et al., 2011). Overall, it appears that the upregulation of HSP70 is often observed in response to cold and freeze stress in insects, but to date this has not been explored in intertidal species.

While molecular chaperones like HSP70 help to mitigate protein damage, freezing can still induce irreversible protein damage and denaturation. In this case, ubiquitin, a low-molecular-mass protein, marks the damaged proteins to be degraded by proteases (Hochstrasser, 1995). Elevated levels of ubiquitin-conjugated proteins accumulating within the cell indicates increased levels of protein damage (Hofmann & Somero, 1995). Elevated levels of ubiquitin conjugates have been observed in cold-adapted notothenioid fish (Todgham et al., 2007), indicating that cold temperatures affect protein integrity at an organismal level. In response to elevated summer temperatures, *M. trossulus* displayed increased levels of ubiquitin conjugates (Hofmann & Somero, 1995). This increase in ubiquitin conjugates was also correlated with increased levels of HSP70, suggesting that chaperone proteins may not sufficiently mitigate protein damage during high temperature stress (Hofmann & Somero, 1995). Taken together, these findings suggest that freezing may induce protein damage, leading to elevated levels of ubiquitin conjugates, and may also induce an increase in molecular chaperone expression.

Another gap in the literature is that the freeze tolerance of intertidal animals has only been investigated through single freezing events, which does not account for the repeated freeze thaw cycles that the animal would experience in nature. While there has been no prior work on the effects of repeated freeze-thaw (RFT) in intertidal species, repeated freezing has several important effects in insects, which are distinct from the effects of single, prolonged freezing events. With repeated freezing events, both a species of hoverfly (Brown et al., 2004) and caterpillar (Marshall & Sinclair, 2011), had decreased survival compared to single sustained freezes of the same length. Similarly freeze thaw cycles induced oxidative stress on the freeze tolerant goldenrod gall fly, and reduced survival compared to a single freeze (Doelling et al., 2014). In response to RFT, goldenrod gall flies also upregulate their sorbitol cryoprotectant levels, but decrease their subsequent egg production (Marshall & Sinclair, 2018). By contrast, there was no significant difference in survival for *B. antarctica* larvae exposed to repeated freezing events compared to those that received a prolonged freeze exposure of equal length (Teets et al., 2011). It is evident that RFT has important effects on some insects, although it is unclear how this may translate to intertidal organisms.

The model organism used in this study is the bay mussel *Mytilus trossulus*, which is a freeze tolerant invertebrate that inhabits rocky intertidal zones along the west coast of North America, as far north as Alaska, and experiences stressful freezing temperatures throughout the winters. When mussels are exposed to the air during low tides, they tightly close their valves, trapping seawater within their shell. When this seawater freezes at low temperatures, this causes the animal to come into direct contact with ice, which induces seed-freezing in the tissues (Storey & Storey, 1996). When mussels freeze completely, over 60% of their total body water can convert to ice (Kanwisher, 1955). Research on cryoprotective qualities of *M. trossulus* and its congener *Mytilus edulis* in response to freeze exposure is limited. Low molecular weight cryoprotectants, like amino acids, are thought to play an important role in preventing osmotic shock in response to freezing in *M. trossulus* (Kennedy et al., 2020). Additionally, the amino acid taurine has been demonstrated to be cryoprotective in *M. edulis* by preserving membrane integrity *in vitro* (Loomis et al., 1988). However, these previous studies were conducted under constant temperature conditions, but fluctuating temperatures (such as RFT) may produce undiscovered effects on *M. trossulus* (Marshall et al., 2021).

There are many unknowns surrounding the physiological and biochemical impacts of freezing; the unknown effects of freeze-thaw cycles on survival compared to prolonged freezing periods and the poorly understood role of HSP70 in mediating protein damage in response to cold and freezing stress. In addition, how an organism’s intertidal shore position and the amount of recovery time an organism has post-freeze impacts these effects is unclear. Here we hypothesize that repeated freeze-thaw cycles are more stressful to mussels, as compared to single, prolonged freezing events. We predict that freeze-thaw cycles will reduce *M. trossulus* survival relative to single freezes, since mussels will need to survive and cope with various stressors during multiple rounds of the cool/freeze/thaw cycle, as opposed to prolonged-freeze-exposed mussels that only experience one round of cool/freeze/thaw. We expect RFT will particularly reduce survival in low shore mussels, since they are not as well acclimated to freeze exposures. We also predict that HSP70 and ubiquitin will be upregulated in response to freeze thaw cycles, as the more strenuous freeze exposures will result in elevated levels of protein damage that will need to be repaired. We expect this trend to be most pronounced in low shore mussels.

## Methods

### Mytilus trossulus *Collection*

Specimens of *Mytilus trossulus* were collected from Tower Beach, Vancouver, British Columbia, Canada (49.2733° N, 123.2578° W) on Oct. 14, Nov. 14, Dec. 13, 2020, and May 25, 2021. Mussel collections were completed using a Scientific License, Management of Contaminated Fisheries Regulations from the Department of Fisheries and Oceans Canada (License number: XMCFR 22 2019). In Vancouver, sea water temperatures usually hover between 7 and 12°C year-round, while winter air temperatures decline as low as −10°C (JPL MUR MEaSUREs Project, 2015).

On each sampling day, mussels were retrieved from the same outcropping of rocks at the lowest daily low tide, which in Vancouver during the winter is at night. High intertidal mussels were located from the uppermost edge of the mussel bed, and low intertidal mussels were collected from just above the water line. Neither sex nor age were controlled for during collections and experiments, though mussels were selected for size which is a proxy for age (Richardson et al., 1990). Animals collected were selected for an approximate size of 2-3 cm. Within 1 hour of collection, mussels were placed in aerated 20 L aquaria in natural seawater (27.5 ppt salinity) sourced from seawater taps in the UBC Zoology Aquatics Facility. The aquaria were placed in 7 °C incubators (MIR-154, Sanyo, Bensenville, USA) in the Biosciences Building at the University of British Columbia, Canada. Approximately 100 mussels were stored in each 10-gallon aquaria. A full seawater change was performed every two days. Once collected, mussels were haphazardly assigned to experimental groups.

### Freeze Exposures

When conducting the freeze exposures, *M. trossulus* were removed from the aquaria, dried with paper towel and labelled with an identification number on the shell with nail polish. Mussel shell length was measured at the longest length of the valves using manual calipers and recorded. Then a copper-constantan type-T thermocouple (OMEGA Engineering, St-Eustache, Quebec) was attached to each mussel’s shell with adhesive putty or small pieces of masking tape. Thermocouples were connected to computers using PicoLog 6 beta software for Windows through Picolog TC-08 interfaces (Pico Technology, Cambridge, UK) throughout the freeze exposure to track mussel body temperatures and record freezing events. This was demonstrated as a rapid release of heat and dramatic rise in body temperature, which happens immediately after the supercooling point is reached (which is the point where body fluids turn to ice; Lee, 2010). *M. trossulus* were stacked in 25 mm *Drosophila* vials, with individuals separated by Styrofoam to avoid inter-individual ice nucleation. The mussels were placed in a cooling bath with an initial temperature of 7 °C (to represent the temperature of the seawater) and then cooled to the desired temperature with a rate of −1 °C/min or −1.5 °C/min (in the survival assay) and −1.5 °C/min for the HSP70 and protein aggregation assays, which roughly mimics the rate of cooling mussels would experience in the field as the tide recedes.

Mussels used in survival trials were collected from the high and low edges of the mussel bed on November 14, 2020 and kept in seawater aquaria at 27.5 ppt salinity for 2-10 days before freezing. Mussels were then frozen at −8 °C for different allotments of time: one continuous exposure for 8 h (1 × 8 h), two exposures that were 4 h each (2 × 4 h) or four separate exposures that were 2 h each (4 × 2 h). −8 °C was chosen as the experimental temperature based on a pilot study that produced 90% mussel survival after a 2 h freeze. After freeze exposure, mussels were taken immediately from freezing baths and placed into seawater aquaria set to 7 °C. Mussels that were repeatedly frozen were allowed to recover in the aquaria for 24 hours between each freeze. After their last freezing exposure, mussels were monitored daily for viability for 7 days, and were determined to be dead if the valves did not close when the animals were removed from the water, or upon mechanical stimuli (light probing with a metal tool).

Mussels used in HSP70 and ubiquitin quantification were collected from the high and low edges of the mussel bed on December 12, 2020 and kept in 25 ppt, 7 °C aerated seawater aquaria for 24 hours until freezing. To investigate how freeze thaw cycles affect HSP70 expression, mussels were frozen once at −6°C for either 8 consecutive hours (1 × 8 h) or four repeated freeze times of 2 hours (4 × 2 h) with 22 hours of recovery between (n=10). −6 °C was chosen as the experimental temperature based on a pilot study done using mussels from this collecting trip that produced 90% mussel survival after a 2-hour freeze. After the final freeze exposure for each treatment group, mussels were returned to their aquaria for 2 hours or 20 hours. The 2 hours allowed for adequate time for protein expression (Miller et al., 2009), and the 20 hour recovery time was used to investigate any possible differences in protein expression with longer recovery times. Additionally, as a control group for time of day, 7 mussels that were not exposed to any freezing treatments were sacrificed at each same timepoint.

Finally, a new subset of *M. trossulus* were collected on May 25, 2021 and stored using the same methods as before but in 17.5 °C seawater (reflecting the temperature of seawater on date of collection). To observe HSP70 expression in *M. trossulus* after a single freeze exposure, mussels were frozen only once at −6°C for 2, 4, 6, or 8 hours and then allowed to recover for 20 hours.

### Sample preparation

After recovery in the aquaria, mussels were removed, and the gill tissue was dissected from the mussels (approximately 100 mg). Gill tissue was chosen over other tissues in the mussel because it has been shown to be the most sensitive to temperature, thus more likely to upregulate HSP70 expression (Aleng et al., 2015). After being blotted with a Kimwipe to remove excess water, gill tissue was weighed and immediately frozen in microcentrifuge tubes at −80 °C and stored until further use. The HSP70 assays were carried out using a protocol similar to that of Sagarin and Somero (2006). Frozen gills were placed in 400 μL of pH 7.1 lysis buffer [32 mM Tris-HCl (pH 7.1), 1 mM Ethylenediaminetetraacetic acid (EDTA), 1 mM sodium dodecyl sulfate (SDS), 0.25 mg/mL phenylmethylsulphonyl fluoride (PMSF), and 10 μg/mL Leupeptin]. Samples were twice boiled in a dry bath for 5 min to denature the proteins, and then homogenized using a bullet blender (Bullet Blender 50 Blue, Next Advance) with 200 μL of 3.2 mm round beads at for 10 min at setting 8 in 1.8 mL microcentrifuge vials (Eppendorf Safe-Lock). Homogenates were transferred to a new microcentrifuge vial and briefly placed on ice for phase separation before centrifugation at 14,000 × *g* for 15 minutes. Supernatant was stored at −80 °C.

Determination of the total protein in each sample was conducted using a BCA Protein Assay Kit (Pierce Inc.) in 200 μL 96 well plates. Each plate contained a set of serially diluted BSA standards, which were loaded in triplicate, and then the samples were also loaded in triplicates. Absorbance of the samples were measured using a spectrophotometer (SpectraMax Plus 384 Microplate Spectrophotometer, Molecular Devices LLC) at a wavelength of 562 nm. A standard curve was generated from the absorbance data, and the average of the three replicates of each sample was calculated to determine the protein concentration. This value was used to determine the volume of each sample to load into the gel wells, since an equal amount of protein is needed to compare HSP70 or ubiquitin expression effectively.

### Western Blots

Protein separation was conducted by electrophoresis in a Mini Gel Tank on 10% Tris-Glycine pre-cast gels (Invitrogen™ Novex™ Value™ 10%, Tris-Glycine, 1.0 mm, Mini Protein Gel, 15-well). In each gel, one lane was dedicated to the prestained marker (Thermo Scientific™ PageRuler™ Plus Prestained Protein Ladder, 10 to 250 kDa), one lane was loaded with 5 μg BSA as a negative control, and another was loaded with 10 ng of human HSP70 (His tagged human HSP70, SRP5190, Millipore Sigma), which allowed comparisons to be made among separate gels. Four μL of NuPAGE™ Reducing Agent (10X) and 20 μL of Tris-Glycine SDS Sample Buffer (2X) were added to 5 μg of protein in the samples and topped up with deionized water to reach a final volume of 40 μL. Samples were then heated to 85°C for 2 minutes. The 13 remaining lanes on the gel were dedicated to these protein samples, and 10 μL of sample was loaded into each well. Each protein sample was loaded 3 times, in separate gels – for a total of 3 technical replicates per protein sample. Loaded gels were submerged in running buffer (Invitrogen™ Novex™ Tris-Glycine SDS Running Buffer) and run for approximately 45 minutes at 225V.

Western blots were used to quantify HSP70 expression. Gels were immediately transferred to 0.45 μm nitrocellulose membranes (BioRad) in a Trans Blot chamber (BioRad) for 6 hours or overnight at 30 V. After equilibration in transfer buffer, gel was sandwiched with nitrocellulose membrane between two pieces of filter paper, and two blotting pads submerged in transfer buffer (Tris-glycine and 20% methanol) during the transfer. After the transfer, the membrane was stained with Ponceau Red for 3 minutes and then destained with water to visualize the protein transfer. Membranes were air dried and stored between filter paper in Ziploc bags at 4°C before the start of the immunoassay.

Immunoassay was conducted using a protocol similar to Miller, Harley, and Denny (2018). Membranes were incubated in blocking buffer [1X Tris-buffered saline (TBS), 3% BSA, 0.1% Tween-20] for 1 hour on an orbital shaker. Membranes were then washed four times in TBS with 0.1% Tween-20 (TBST) for 5 minutes each. Then, membranes were incubated in a 1:5000 dilution of the primary antibody in PBS with 2.5% BSA (antibody MA3-007, clone 5A5, Mouse Monoclonal Antibody, FisherScientific). The membranes were washed four times in TBST for 5 minutes each before incubation in secondary antibody (A1418, Anti-mouse IgG (Fc specific) alkaline phosphatase antibody produced in goat, Sigma-Aldrich). The secondary antibody was diluted 1:1000 in TBST with 2.5% BSA, and the membrane was incubated for 45 minutes. The membranes were then washed four times for 5 minutes each in TBST.

The proteins were visualized by incubating membranes for 30 minutes in BCIP®/NBT Alkaline Phosphatase Substrate (B5655, SIGMAFAST™ BCIP®/NBT, Sigma-Aldrich). The intensities of the detected bands were analyzed with a FluorChem 8800 Imager (Alpha Innotech, San Leandro, CA, USA) using AlphaEase FC software (v. 3.1.2; Alpha Innotech). The density measurements of each protein sample band at 68, 72 and 76 kDa, were compared relative to the human HSP70 standard on each gel, allowing for comparison of relative density values across all western blots.

### Dot Blots

Using the same protein samples as with the HSP70 assay, ubiquitin-conjugated protein expression was examined using immunochemical analysis. 0.2 μm nitrocellulose membranes were prewet for 10 minutes in TBS then secured into the Bio-Dot Microfiltration apparatus (Bio-Rad). Wells were rehydrated once with TBS, then 0.5 μg of each protein sample and 0.05 μg of ubiquitin standard (AB218616, Recombinant Human Ubiquitin protein (His tag), Abcam) were blotted in triplicate and allowed to gravity filter. Wells were washed twice with 200μL TBST using vacuum filtration and the membrane was removed from the apparatus and heat fixed at 55 °C for 20 minutes. Membranes were then developed and visualized using the same protocol as the HSP70 assay, except for an ubiquitin specific primary antibody (1:1000 dilution; UBCJ2, Mono- and polyubiquinated conjugates monoclonal antibody, FroggaBio).

### Statistical analysis

Statistical analyses were performed using R (c. 3.5.1; R Development Core Team, 2013). The R package “ggplot2” was used to generate figures 1 and 3-8 (Wickham, 2016), the package “cowplot” was also used to generate Figure 1, 3, 4, 5, 7, 8 (Wilke, 2020). The R package “plyr” was also used to determine means and standard errors before plotting (Wickham, 2011).

**Figure 1:**
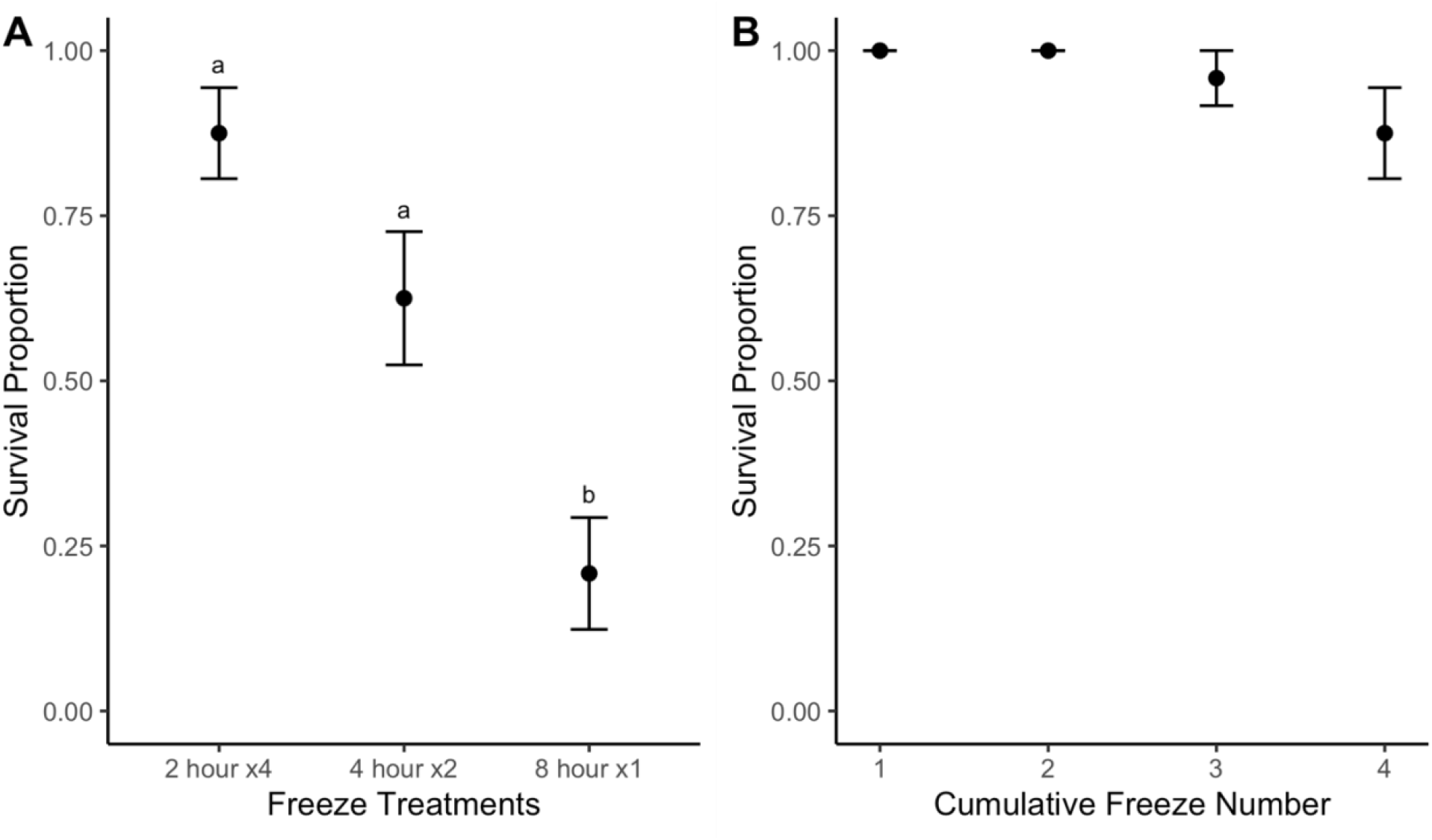
A) Bay mussels (M. trossulus) are more tolerant to repeated freeze-thaw cycles rather than a single, sustained freeze B) Each 2 h freeze-thaw cycle induces only a small, non-significant, increase in mortality in M. trossulus. Mussels were collected from Tower Beach on Dec 12, 2020 and were exposed at −8 °C for either 4 × 2 h, 2 × 4h or 1 × 8 h, with a cooling rate of −1 or −1.5 °C/minute (n=12 per treatment). Mussels were allowed to recover in 25 ppt, 7°C seawater for 24 hours between each freeze. Survival was assessed for 7 days following freeze exposure. Error bars are standard error of the proportion.

Logistic regression was used to test for significant differences in survival within different freeze-thaw cycle lengths. Then, a chi-squared test was used to test for significant differences in survival after freezing assessing shell length, total time frozen, and shore height as potential predictors of survival after freezing. Specific differences between survival groups were analyzed using an ANOVA and Tukey HSD post-hoc test (Tukey, 1977). To determine how basal levels of HSP70 changed with shore height, HSP70 density values were standardized to the HSP70 human standard (so they could be accurately compared among separate blots) and then tested for normality using a Shapiro Wilk test (Royston, 1982). If data was normal, the means of the shore height were compared using a student’s t-test, and if it was not normal, a Mann-Whitney U (Mann & Whitney, 1947) test was used. Finally, freeze time, recovery and shore position were analyzed in relation to HSP70 density. First, relative HSP70 density was calculated by taking the densitometry measurements of the sample and dividing them against the densitometry value for the HSP70 human standard, and then by the means of their time matched controls. The resulting values were then log transformed (expressed as log_2_ fold change) to better adjust for normality. A three-way ANOVA was used to determine if freeze time, recovery, or shore position had an effect on HSP70 density, as well as recording interaction terms between the factors. The same statistical tests were used to evaluate ubiquitin levels in relation to shore height, freeze time, and recovery time. Values are reported as means ± standard error. Alpha was set to 0.05.

## Results

### Effects of Repeated Freeze Thaw on Mussel Survival

To test the effects of repeated freeze thaw cycles on mussels, 168 mussels were frozen either for 1 × 8 h, 2 × 4 h or 4 × 2 h. On average, mussels froze at −3.3 ± 0.15°C (n=60), and every mussel in this experiment froze, as evidenced by the characteristic exotherm in the temperature trace graph that follows the supercooling point. There was no significant effect of shore position within the intertidal zone on survival (df= 1, 68, z=1.01, p= 0.15), and shell length was also not a significant predictor of survival (df= 1, 67, F=3.5, p=0.06). However, mussels that were repeatedly frozen for 4 × 2 h (proportion = 0.88) or 2 × 4 h (proportion=0.61) had higher survival than mussels that had a single prolonged freeze exposure (proportion=0.21; df=1,70, p<0.0001; Figure 1A). Freezing twice for 4 h each time caused similar mortality as freezing four times for 2 h each (p=0.106; Figure 1A). Cumulative survival in *M. trossulus* repeatedly frozen for two hours stayed constant for the first two freezes, but then decreased slightly with more freeze–thaw cycles, but this trend was not statistically significant (Z=-1.76, p= 0.08; Figure 1B).

### Relative HSP70 Expression Analysis via Western Blot

A new subset of 88 mussels was collected on Dec 13^th^, 2020 and frozen for either 4 × 2 h or 1 × 8 h at −6 °C, and all but 1 survived until their dissection time. Mussels froze at −4.4 ± 0.8°C (n=136), and every mussel in this experiment froze.

HSP70 expression was determined through semi-quantitative densitometry analysis from Western blots. The monoclonal antibody used in the immunoblot analysis (FisherScientific, MA3-007) recognized three different isoforms of HSP70 (at 68, 70 and 76 kDa) in the gill tissue of *M. trossulus*. In some specimens, the bottom two bottom bands were not well resolved, therefore comparisons of HSP70 expression were performed using either all bands, the top band (76 kDa) or the bottom two bands (68+70 kDa). A representative western blot is shown in Figure 2.

**Figure 2:**
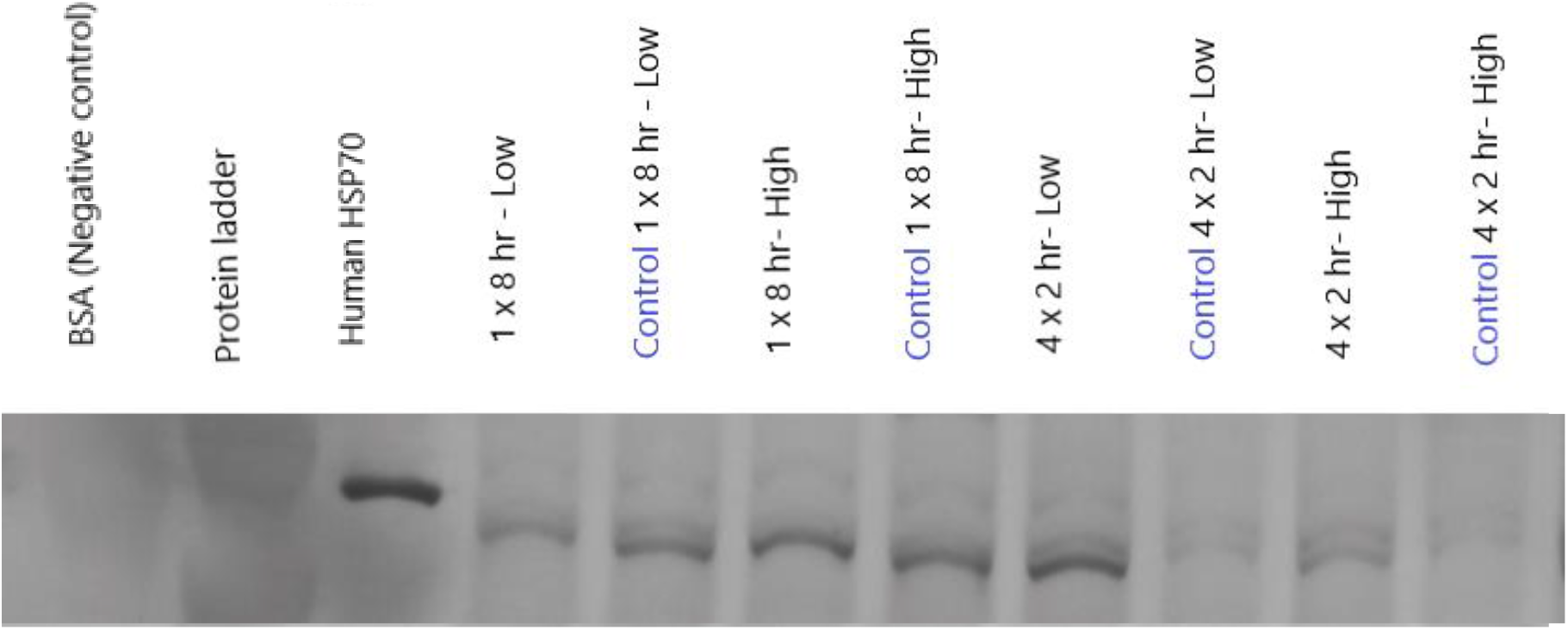
Western blot analysis of gill tissue HSP70 expression from mussels frozen at −6°C. Equivalent amounts of gill protein (5 μg) were separated on a 10% Tris-glycine gel, followed by electrotransfer and immunostaining using alkaline phosphatase antibodies. Labels indicate the treatment group or time-matched controls, and all treatment groups on this blot were allowed to recover for 20 h. The standard was 10 ng of human HSP70.

Based on semi-quantitative densitometry, concentration of the 76 kDa isoform was significantly higher in gills of mussels from the high intertidal zone compared to the low intertidal zone before any freezing event occurred (t = 4.0, p < 0.001; Figure 3C). Despite the significant difference in the 76 kDa isoform, this difference was not observed in the bottom 2 bands (p = 0.74; Figure 3B) or when all bands were combined (p = 0.32; Figure 3A).

**Figure 3:**
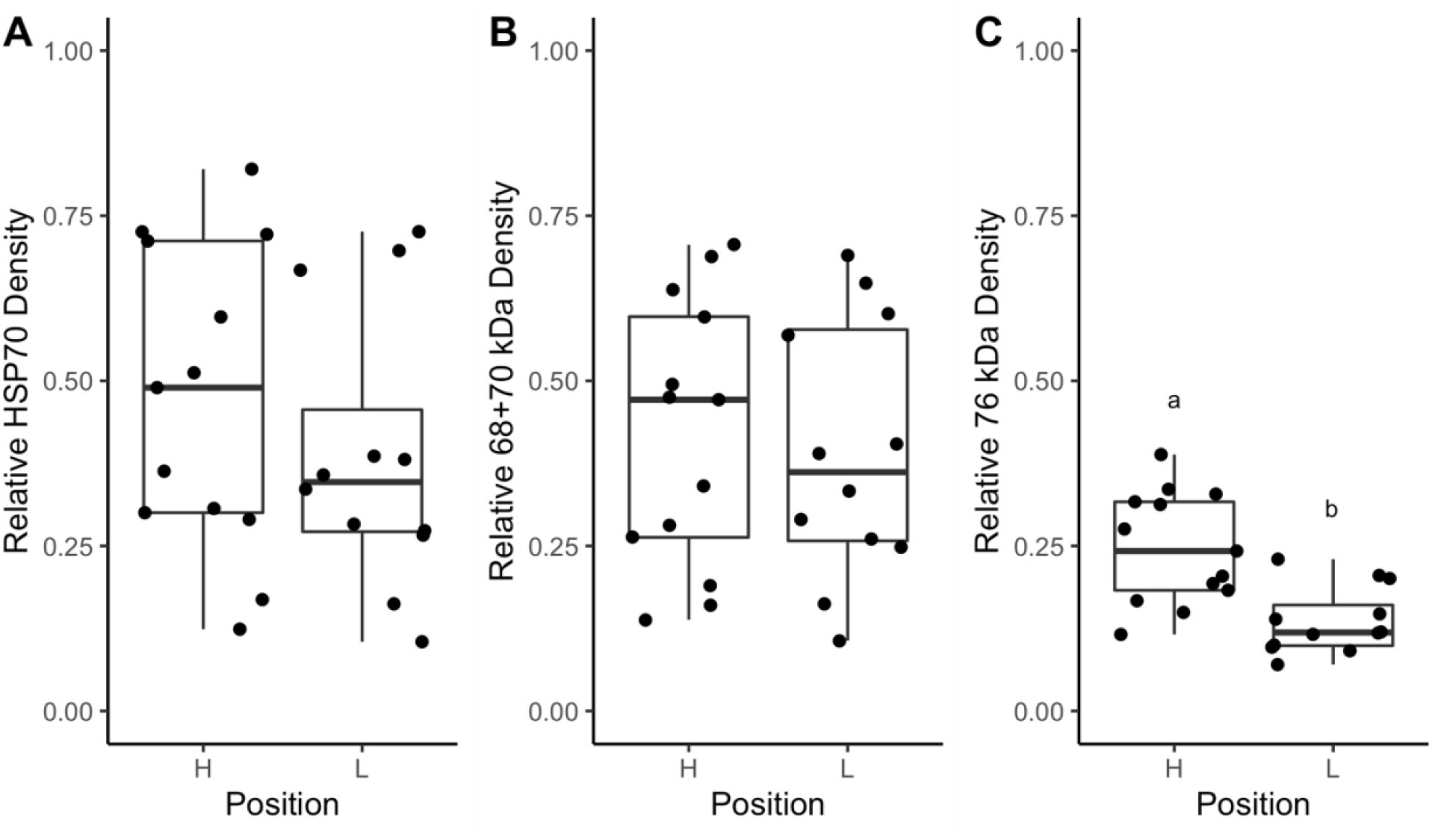
High intertidal M. trossulus have higher basal expression of the 76 kDa isoform, as compared to low intertidal mussels. A) Relative HSP70 expression of all bands. B) Relative HSP70 expression of bottom bands (68+70 kDa). C) Relative HSP70 expression of top bands (76 kDa). Mussels were collected at Tower Beach, Vancouver on Dec 12^th^ 2020, and were not exposed to any freezing. Horizontal lines represent the medians, boxes represent the interquartile range, and lower and upper error lines show 10th and 90th percentile, respectively. Overlain points represent the HSP70 density for each individual mussel gill tissue sample, the asterisk indicates statistical significance. Densities are relative to HSP70 human standard. n=13 for high, and n=12 for low.

### HSP70 expression in response to repeated freeze-thaw

Each frozen mussel was directly paired with a control mussel that remained in seawater that was dissected at the same time to reduce any circadian effects (Figure 1). Therefore, the densitometries have been displayed as HSP70 expression relative to the average of the time-matched controls (expressed as log_2_ fold change).

Neither freeze treatment or recovery time had a significant effect on relative HSP70 density for the sum of the 68+70 kDa isoforms (Freeze Treatment: F_(1, 24)_ = 0.13 p= 0.72; Recovery: F_(1, 24)_ = 0.96 p= 0.34) or the 76 kDa isoform (Freeze Treatment: F_(1, 24)_ = 0.02 p= 0.88; Recovery: F_(1, 24)_ = 1.0 p= 0.32). When all HSP70 isoforms were combined, recovery time had a significant effect on HSP70 density, but not freeze treatment (Freeze Treatment: F_(1, 32)_ = 2.8 p= 0.1; Recovery time: F_(2, 32)_ = 4.3 p= 0.045). In mussels exposed to repeated freeze thaw cycles, average density of total HSP70 protein within gill tissue was greater following a 20-hour recovery (Freeze Treatment × Recovery: F_(1, 24)_ = 5.2 p= 0.03; Fig. 4A). Expression of HSP70 in the bottom two bands (F_(1, 24)_ =1.6, p= 0.21; Fig. 4B) and the top band isoform (F_(1, 24)_ = 1.9, p= 0.18; Fig. 4C) did not have a significant interaction term between recovery times and freeze treatment.

**Figure 4:**
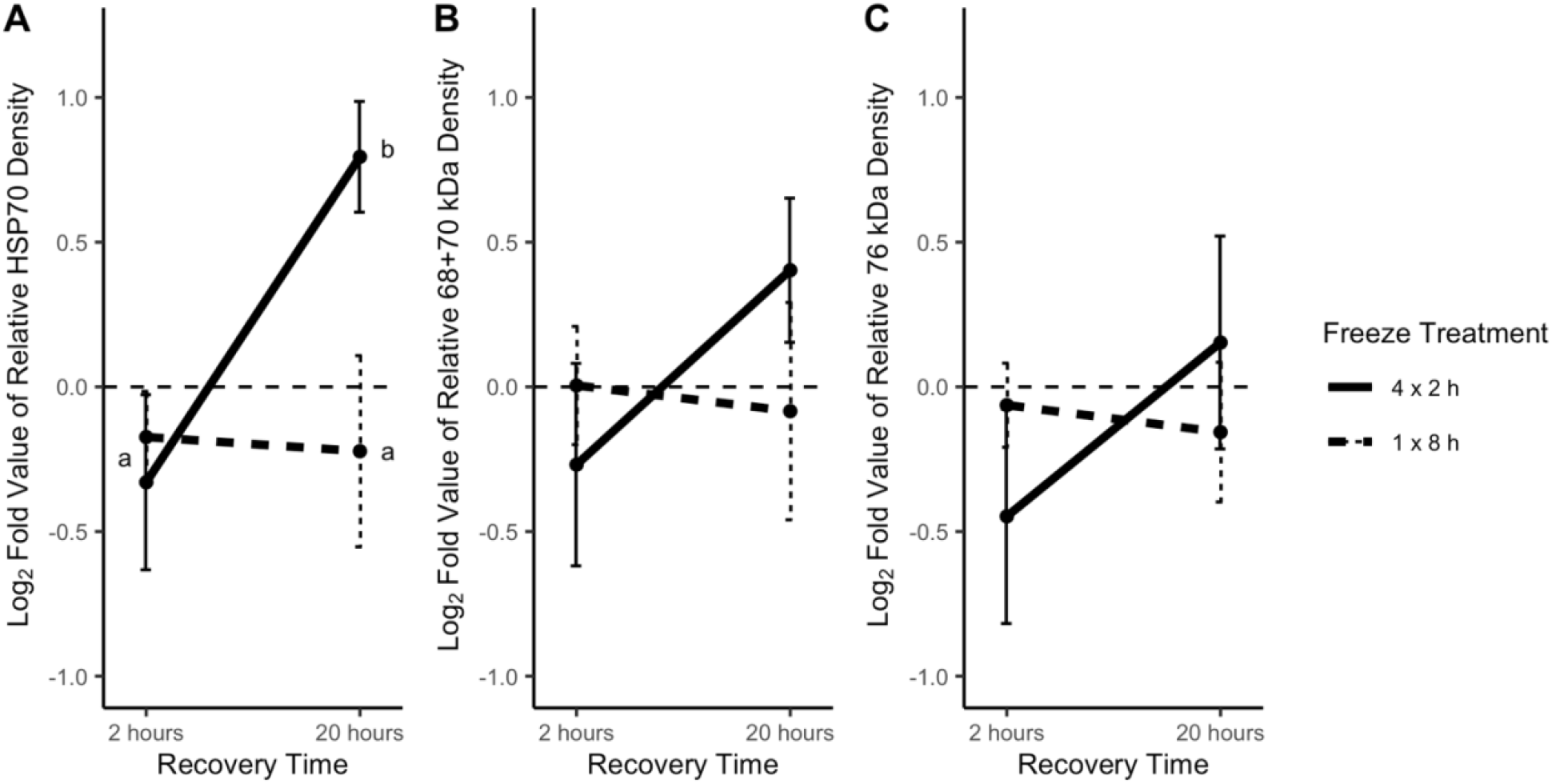
Relative HSP70 density significantly increases in M. trossulus only after repeated freeze-thaws, and only after 20 h recovery. A) Relative log fold change HSP70 expression of all bands. B) Relative log fold change HSP70 expression of bottom bands (68+70 kDa). C) Relative log fold change HSP70 expression of top bands (76kDa). Measurements are relative to 10 ng human HSP70 standard, and then expressed as log2 fold change relative to time matched control. Mussels in the 4 × 2 h freeze treatment group were frozen four times for 2 hours at −6 °C, with a recovery period of 22 hours in between each freeze. Mussels in the 1 × 8 h freeze treatment group were frozen once for 8 hours. Letters ‘a’ and ‘b’ represent statistical significance. Dashed line represents 0, or no change in relative HSP70 expression relative to time-matched control. Error bars represent standard error of the mean. n=8.

There was a significant interaction between shore position and freeze treatment such that only mussels from the low intertidal exhibited increased relative 76 kDa isoform expression following repeated freeze thaw cycles (Position: F_(1, 28)_ =8.4 p<0.01; Freeze Time × Position: F_(1, 28)_ =6.1 p=0.02; Fig. 5C). When exposed to repeated freeze-thaw cycles, low intertidal mussels had a higher expression of the 76 kDa isoform (mean = 0.53 ± 0.19) than their high intertidal counterparts (mean = −0.82 ± 0.36; Fig. 5C). Repeated freeze-thaw cycles induced HSP70 expression in low intertidal mussels compared to control mussels, in all bands (mean = 0.58 ± 0.31), bottom bands (mean = 0.47 ± 0.25) and top band (mean = 0.53 ± 0.19; Figure 5).

**Figure 5:**
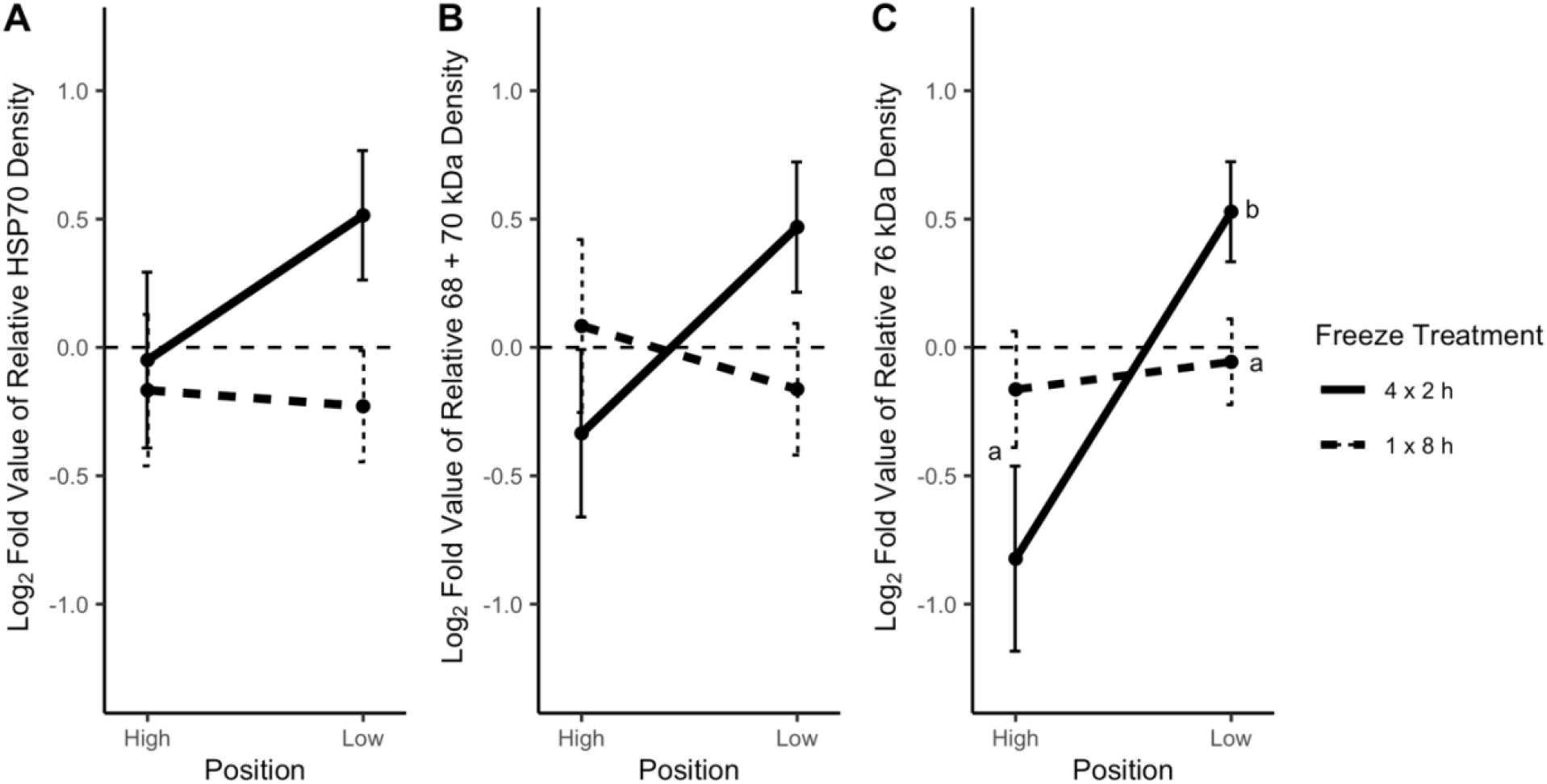
Repeated freezing only induced HSP70 in low intertidal M. trossulus. A) Relative log fold change HSP70 expression of all bands. B) Relative log fold change HSP70 expression of bottom bands (68+70 kDa). C) Relative log fold change HSP70 expression of top bands (76 kDa). Measurements are relative to 10 ng human HSP70 standard, and then expressed as log2 fold change relative to time matched control. Mussels in the 4 × 2 hour freeze treatment group were frozen four times for 2 hours at −6 °C, with a recovery period of 22 hours in between each freeze. Mussels in the 1 × 8 hour freeze treatment group were frozen once for 8 hours. Letters ‘a’ and ‘b’ represent statistical significance. Dashed line represents 0, or no change in HSP70 expression relative to time-matched control. Error bars represent standard error. n=8.

There was no effect of shore position on HSP70 expression in the mussels that experienced a sustained freezing event (1 × 8 h) in any of the isoforms (p > 0.09 in all cases; Figure 5).

To determine if mussels were able to upregulate HSP70 in response to a singular freeze event of any length, *M. trossulus* were frozen for a single freeze of either 2, 4, 6, or 8 hours. No significant upregulation in HSP70 expression was observed among any of the freeze times (p > 0.3; Figure 6).

**Figure 6:**
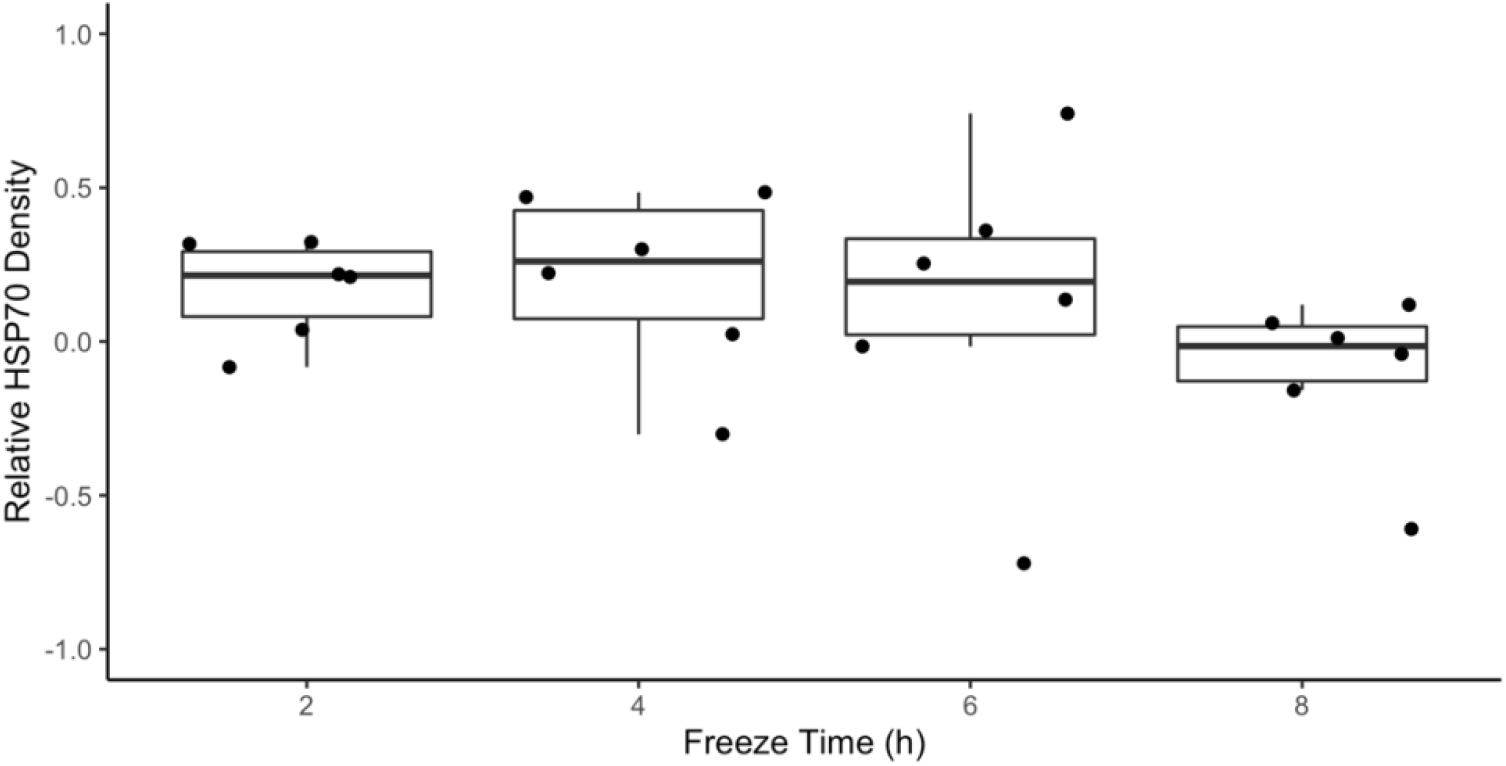
Singular freezes of M. trossulus at −6 °C. Relative HSP70 expression using all bands at 2, 4, 6, and 8 hours of freezing at −6°C, at a cooling rate of −1.5 °C/minute. Mussels were allowed to recover for 20 hours prior to HSP70 analysis. Horizontal lines represent the medians, boxes represent the interquartile range, and lower and upper error lines show 10th and 90th percentile, respectively. Overlain points represent the HSP70 density for each individual mussel gill tissue sample. Measurements are relative to human HSP70 standard, and then expressed as log2 fold change relative to time matched control. n = 6.

### Relative Ubiquitin Expression Analysis via Dot Blot

Using the same protein samples as in the HSP70 analysis, ubiquitin expression was analyzed using dot blotting. Basal ubiquitin expression did not differ between *M. trossulus* from the high intertidal and mussels from the low intertidal (p > 0.35; Figure S1). When exposed to freeze-thaw cycles, mussels that recovered for 20 hours had a significantly higher ubiquitin expression (mean = 0.06 ± 0.05) than those that recovered for 2 hours after freeze exposure (mean = −0.16 ± 0.03; p < 0.005; Fig. 7A). There was no significant effect of shore position on ubiquitin expression (p=0.45; Figure 7B).

**Figure 7:**
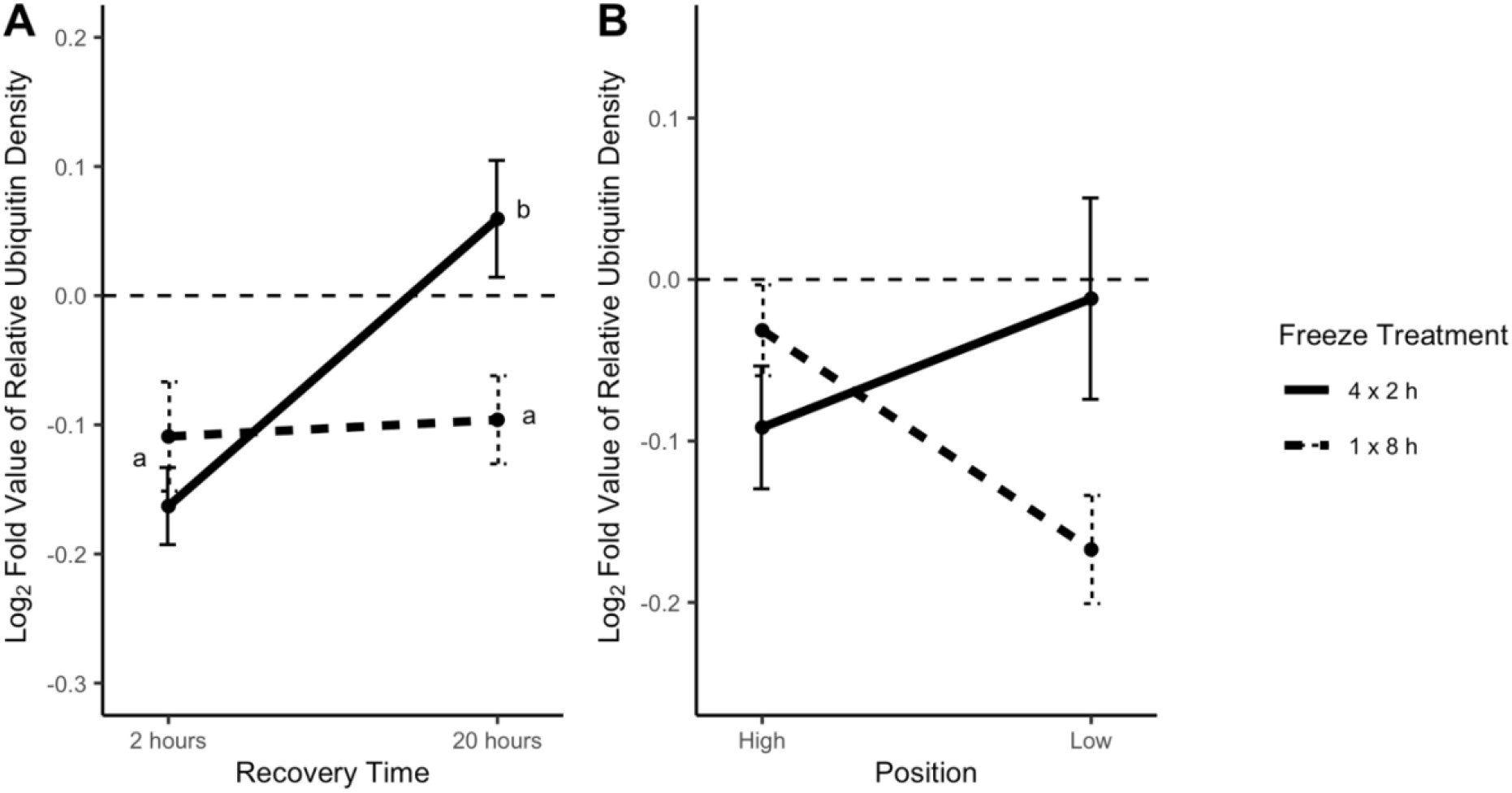
20 hours of recovery after repeated freezing increases relative ubiquitin density increases in M. trossulus, with no significant effect on the shore height of the mussels. A) Relative log2 fold change of ubiquitin expression after 2 and 20 hours of recovery post-freezing. B) Relative log fold change of ubiquitin expression from high and low intertidal mussels. Measurements are relative to 50 ng of ubiquitin standard, and then to time matched control. Mussels in the 4 × 2 h freeze treatment group were frozen four times for 2 hours at −6 °C, with a recovery period of 22 hours in between each freeze. Mussels in the 1 × 8 h freeze treatment group were frozen once for 8 hours. Dashed line represents 0, or no change in relative HSP70 expression. Error bars represent standard error. n=10.

### HSP70 and ubiquitin correlation

Relative ubiquitin levels and relative HSP70 levels are positively correlated when looking at treatment group averages (R = 0.85; p < 0.01) and when looking at each sample individually (R = 0.46; p <0.05; Figure 8).

**Figure 8:**
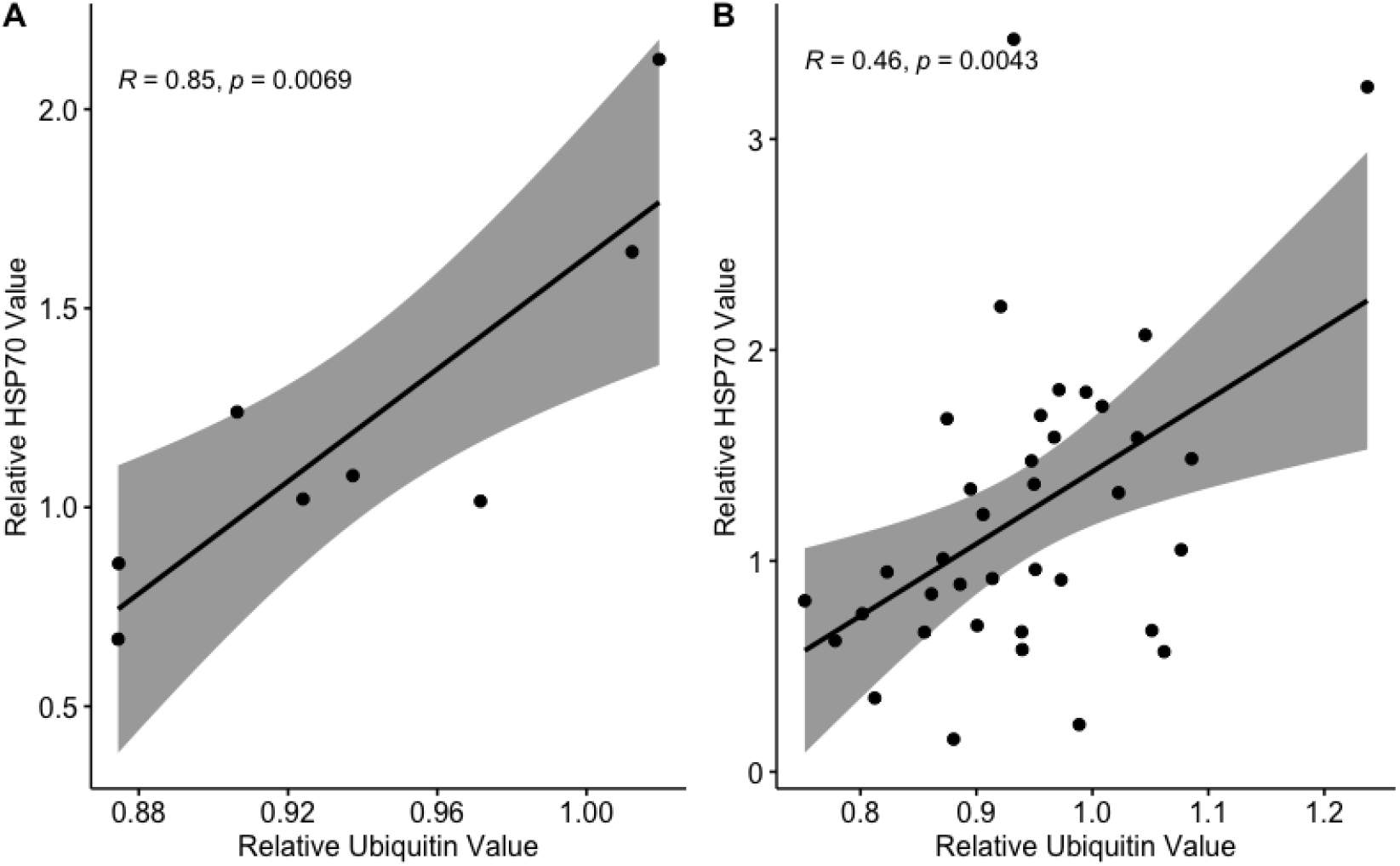
HSP70 expression and ubiquitin levels are positively correlated in M. trossulus after freeze exposures. A) Correlation between relative HSP70 values and relative ubiquitin values averaging the individual samples from each treatment B) Correlation between relative HSP70 values and relative ubiquitin values for each individual sample. HSP70 measurements are relative to 10 ng of human HSP70, and ubiquitin measurements are relative to 50 ng of ubiquitin standard. Grey shading represents 95% confidence interval. n=37

## Discussion

Here we show for the first time that HSP70 and ubiquitin are upregulated in response to repeated freezing in *M. trossulus.* HSP70 expression and levels of ubiquitin conjugates in repeatedly frozen *M. trossulus* were investigated to better understand changes to the cellular protein pool during freezing. HSP70 expression was measured by Western blot in mussels that had either experienced repeated freezing or a single sustained freeze. Ubiquitin conjugated proteins were also measured through dot blot analysis for both repeatedly and prolonged frozen mussels. The impacts of shore height of the mussels and the time given for repair was also examined. Overall, we had three salient findings: 1) survival of repeatedly frozen *M. trossulus* was significantly higher than mussels that were frozen only once, 2) basal levels of HSP70 were reduced pre-freezing but then significantly upregulated post-freezing in low intertidal mussels, as compared to high intertidal mussels 3) both HSP70 expression and levels of ubiquitin conjugates were upregulated 20 hours after repeated freezing. Taken together, this may indicate that repair processes during immersion are important for mussels to survive sub-zero air temperatures.

### Survival

*Mytilus trossulus* living on polar and temperate shores can experience many freeze-thaw cycles throughout the winter, and therefore must respond to cellular damage from freezing to survive. Generally, to minimize the effects of freeze-induced damage, animals can either upregulate protective mechanisms before and during freezing, or they can repair their cellular damage after the freeze exposure. Given that animals can respond biochemically to a freezing event, this suggests that the timing and repetition of freezing events might be important for predicting survival. Sustained freezing in *M. trossulus* induced more mortality than repeated freezing (Figure 1), even though the cumulative amount of time spent frozen was held constant. This suggests that during RFT, either cellular repair occurs during each recovery period to mitigate damage, and/or that mussels can upregulate cryoprotective defense mechanisms between each freeze.

While this is the first study to examine the effects of freeze-thaw cycles in intertidal species, there have been previous studies examining the effects of RFT in insects, which have yielded contradictory results about the impact of RFT on survival (Marshall & Sinclair, 2012). RFT resulted in increased mortality for insect species like the goldenrod gall fly *Eurosta solidaginis* (Doelling et al., 2014), woolly bear caterpillar *Pyrrharctia isabella* (Marshall & Sinclair, 2011), and hoverfly *Syrphus ribesii* (Brown et al., 2004). However, *Belgica antarctica* midge larvae had increased survival after RFT compared to a single, sustained freeze (Teets et al., 2011). The findings from our study reveal that *M. trossulus* exposed to RFT have significantly increased survival compared to those that experienced prolonged freeze exposures. During recovery, *M. trossulus* may recover, repair damage, and upregulate defensive mechanisms in preparation for the next freezing event. Perhaps since intertidal animals experience frequent thawing periods in their natural environment in accordance with daily submersion during high tide, they are better able to utilize these “recovery periods” to repair damage after freezing, as compared to insects which aren’t exposed to frequent, predictable thawing periods and are therefore more accustomed to sustained bouts of freezing.

### Intertidal Shore Height Effects

Mussels living higher on the shore are exposed to aerial conditions for longer, and therefore must contend with greater thermal stress compared to their low intertidal counterparts (Halpin et al., 2002). This suggests that mussels from the high edges of the mussel bed may be better adapted to frequent freeze thaw cycles. However, our results indicated no difference in overall survival between high and low intertidal mussels when exposed to freezing events of any duration. This contrasts with Kennedy et al. (2020)’s findings that showed *M. trossulus* from the high intertidal zone survived significantly better than ones from the low intertidal zone during three-hour freeze exposures. The differences found by Kennedy et al. (2020) were relatively small, however, and since the authors of that study used multiple test temperatures, we might not have been picked up on these small differences since we only used one test temperature in our study.

Aside from survival, intertidal organisms from high and low shore positions may also have differences in constitutively expressed cryoprotectants. The difference in *M. trossulus* survival due to shore height is likely not driven by low molecular weight cryoprotectants (Kennedy et al., 2020), which leaves high molecular weight cryoprotectants such as ice nucleating agents, heat shock proteins, or antifreeze proteins to explain these differences. Despite finding no difference in survival proportion, our results showed that *M. trossulus* from the high intertidal zone had a greater basal expression of the 76 kDa heat shock protein compared to mussels from the low intertidal zone. The 76 kDa isoform of HSP70 in *M. trossulus* has been previously identified to be less heat-inducible (Hofmann & Somero, 1995), and combined with our findings that its endogenous expression varies based on shore position, this suggests that the 76 kDa isoform is constitutively expressed in *M. trossulus*. Past research has similarly found that HSP expression can vary with shore height, as *M. trossulus* from a more stressful intertidal habitat had higher basal HSP70 expression compared to a subtidal habitat (Hofmann & Somero, 1995)*. M. californianus* and a limpet species of the genus *Lottia* from high intertidal sites also had higher HSP70 expression, compared to their low intertidal counterparts (Dong et al., 2008; Halpin et al., 2002). In summary, animals inhabiting the high intertidal zone may have higher expression of constitutive HSPs as a preparative and/or protective function for more frequent and extreme thermal stress.

Aside from constitutively expressed cryoprotectants, intertidal organisms from high and low shore positions may differ in the expression of cryoprotective molecules during and after freezing. Our results showed that low intertidal mussels exposed to freeze thaw cycles induced greater expression of HSP70 compared to high intertidal mussels. This may suggest that freeze thaw cycles are more stressful for low intertidal mussels, and because of their lower levels of constitutive cryoprotectants, they must induce higher levels of HSP70 during and/or after freezing to mitigate protein damage. High intertidal mussels, on the other hand, may express greater basal levels of high molecular weight cryoprotectants, like HSP70, to pre-emptively combat the more frequent and extreme thermal stress they experience *in situ*.

### HSP70 expression and levels of ubiquitin conjugates

HSP70 was upregulated in *M. trossulus* in response to freeze thaw cycles only later in recovery. This response was only observed when mussels had been repeatedly frozen, not after a single 2 h or 8 h sustained freeze, and only 20 hours post freezing. *M. trossulus* also upregulated ubiquitin conjugates after being repeatedly frozen. Similarly, ubiquitin conjugates were only upregulated 20 h after the last repeated freezing event. This may come as a result of two separate scenarios: 1) repair of cellular damage after freezing or 2) upregulation of protective mechanisms after freezing to prevent further damage from subsequent freezing events.

In this case, *M. trossulus* most likely upregulated HSP70 and ubiquitin after freeze thaw cycles to repair damage. This is because HSP70 was uniquely upregulated after RFT, which means that RFT either uniquely induces damage, or that 8 hours of freezing induces such severe damage, that *M. trossulus*’ ability to repair freezing-induced damage is extremely hindered. Evidence points to the latter, because mussels that were frozen for 8 hours had significantly higher mortality. Another key finding was that HSP70 peaks in its expression at 20 hours post freezing but is not significantly different at the 2 hour mark. The closely related congener, *M. galloprovincialis*, similarly exhibits peak HSP70 expression 15 hours following a single heat shock, indicating that it takes some time post-thermal stress for significant upregulation of HSP70 to occur in mussels (Cellura et al., 2006). Depending on the type of tide cycle, 20 hours may be enough time for mussels to repair freezing damage before the next low tide exposure that may bring another freezing event. Overall, the findings of this study suggest that *M. trossulus* elevates HSP70 after repeated freezing, likely to repair protein denaturation due to freezing.

Alternatively, this observed effect could be due to HSP70 being upregulated as a way to prevent further freeze induced damage, in preparation for potential subsequent freezing exposures. The freezing events in this experiment were separated by 22 hours in seawater, loosely modelling the immersion/emersion times of a low intertidal mussel experiencing a mixed semi-diurnal tidal cycle with one aerial exposure per day. For this reason, it is a possibility that the elevated expression of HSP70 in *M. trossulus* shortly before the next freezing event serves a preparative or protective function in animals exposed to repeated freezing.

The upregulation of ubiquitin 20 hours after RFT also gives support to the repair scenario, through a few possible mechanisms. The number of ubiquitin-conjugated proteins may directly reflect the number of denatured proteins within the cell (Hofmann & Somero, 1995). If this is the case, we can infer that RFT induces more protein damage compared to sustained freezes, or that 8 hour sustained freezes are excessively damaging, in that *M. trossulus* was not able to tag denatured proteins with Ub. This response may also be a result of disrupted proteostasis within cells, causing a more complex chain of reactions due to freezing. For example, proteins that degrade Ub-tagged molecules, called proteases, could exhibit reduced activity due to freezing stress (Todgham et al., 2007). This would result in a build-up of Ub-conjugates within the cell over the thawing period, as we observed in *M. trossulus* that had been repeatedly frozen. In either of these cases, ubiquitin molecules seem to play a role in repairing mussel cells’ disrupted protein environment after repeated freeze thaw.

### Concluding Remarks

In summary, freeze-thaw cycles result in higher survival in *M. trossulus* compared to sustained freezes of the same total duration. Additionally, after repeated freeze thaw cycles, but not after sustained freezing, mussels upregulate HSP70 and ubiquitin. This likely indicates that freeze-thaw cycles, which happen naturally in the intertidal, provide mussels with a period to repair damage and are therefore crucial for *M. trossulus* survival in sub-zero temperatures. Future studies should examine the time course of HSP70 and ubiquitin expression throughout and after freezing, to see when elevated expression starts, peaks, and ends. Further investigations into other freeze tolerant intertidal species would also be beneficial to understand if other intertidal species modify HSP70 and/or ubiquitin levels in response to freezing in the same way as *M. trossulus*.

## Supporting information

Supplemental results

## Acknowledgements

K.E.M. is supported by an NSERC Discovery grant, J.R.K. is supported by a NSERC CGS-M, and L.T.G. was supported by a NSERC USRA during this work. The authors thank Drs. Christopher Harley and Patricia Schulte for their assistance and ideas which improved this manuscript greatly.

