## Supplemental results for "Proteostasis in ice: The role of heat shock proteins and ubiquitin in the freeze tolerance of the intertidal mussel, *Mytilus trossulus*"

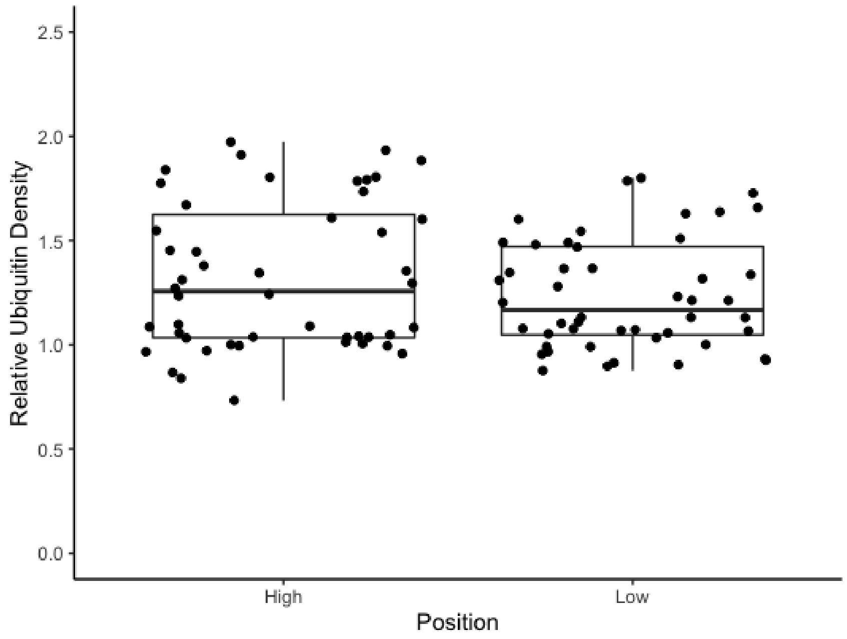


***Figure S1:*** **M. trossulus** ***from the high intertidal and low intertidal have no significant difference in basal ubiquitin expression.*** *Mussels were collected at Tower Beach, Vancouver on Dec 12$^{th}$ and were not exposed to any freeze exposure. Horizontal lines represent the medians, boxes represent the interquartile range, and lower and upper error lines show 10th and 90th percentile, respectively. Overlain points represent the HSP70 density for each individual mussel gill tissue sample. Relative density was calculated by taking the densitometry measurements of the sample and dividing them against the densitometry value for the ubiquitin standard. n=13 for High, and n=12 for Low.*
